# Mechanosensor Piezo1 Mediates Bimodal Patterns of Intracellular Signaling

**DOI:** 10.1101/2022.05.19.492730

**Authors:** Yijia Pan, Linda Zhixia Shi, Daryl Preece, Veronica Gomez-Godinez, Chi Woo Yoon, Shaoying Lu, Christopher Carmona, Seung-Hyun Woo, Shu Chien, Michael W. Berns, Longwei Liu, Yingxiao Wang

## Abstract

Piezo1 belongs to mechano-activatable cation channels serving as biological force sensors. However, the molecular events downstream of Piezo1 activation remain unclear. In this study, we used biosensors based on fluorescence resonance energy transfer (FRET) to investigate the dynamic modes of Piezo1-mediated signaling and revealed a bimodal pattern of Piezo1-induced intracellular calcium signaling. Laser-induced shockwaves (LIS) and its associated shear stress can mechanically activate Piezo1 to induce transient intracellular calcium (Ca_[i]_) elevation, accompanied by an increase in FAK activity. Interestingly, multiple pulses of shockwave stimulation caused a more sustained calcium increase and a decrease of FAK activity. Similarly, tuning the degree of Piezo1 activation by titrating either the dosage of Piezo1 ligand Yoda1 or the expression level of Piezo1, produced a similar bimodal pattern of FAK responses. Specifically, a low degree of Piezo1 activation (transient mode) leads to a transient Ca_[i]_ response with FAK activation, whereas a high degree of Piezo1 activation (sustained mode) causes a sustained Ca_[i]_ response with FAK suppression. Further investigation revealed that SHP2 serves as an intermediate regulator mediating this bimodal pattern in Piezo1 sensing and signaling. These results suggest that the degrees of Piezo1 activation induced by both mechanical LIS and chemical ligand stimulation may determine downstream signaling characteristics.

## 1. Introduction

Cells constantly encounter mechanical forces in the forms of shear stress, compression and stretching. Mechanotransduction is crucial for cells to perceive the local microenvironments in order to maintain cellular function and tissue homeostasis ^1-4^. Alterations of mechanotransduction can lead to pathological conditions ^5^. However, it remains unclear on the detailed mechanism by which cells perceive mechanical cues of the external microenvironment and transduce them into molecular signals and genetic regulations for the coordination of cellular responses.

Several families of molecular mechanoreceptors have been reported to transduce mechanical stimulation into ion currents to modulate physiological functions, particularly the transmembrane ion channel proteins ^6^. Piezo1 has been identified as a component of mechanically activated cation channels in both vertebrates and non-vertebrates. It induces currents in various cell types upon the application of mechanical pressure, tension, or deformation to the cell membrane ^7-9^. Most of the mechanical stimulations resulted in transient calcium influx, although sustained calcium influx upon prolonged mechanical stretch was also reported ^10^. Consequently, Piezo1 plays different roles in physiology, including angiogenesis, vasculogenesis ^11^, and erythrocyte volume regulation ^12^. Piezo1 is also involved in regulating cancer biology in complicated manners. For instance, blocking Piezo1 decreased the motility of the breast cancer cell line MCF-7 ^13^, but increased the migration speed of small cell lung cancer cell lines ^14^. These results suggest a complex role of Piezo1 in regulating cancer cell migration under different biological and environmental contexts.

Due to the important role of Piezo1 in regulating physiological and pathological cell migration, an emerging topic of interest has been focused on the dynamics and effects of Piezo1 activation on cytoskeleton^15^. Focal adhesion zones are hubs for cytoskeletal structures that connect the extracellular matrix to the intracellular cytoskeleton, and they can detect and transmit external mechanical forces. Focal adhesion kinase (FAK) belongs to focal adhesion zone proteins and is a key regulator of focal adhesion dynamics and cytoskeletal remodeling. Therefore, studying the effect of Piezo1 activation on FAK dynamics can give insights into how Piezo1 affects cellular migration. In this study, we applied laser-induced shockwaves (LIS) and chemical stimulations to activate the Piezo1 in live cells and studied the mechanism by which Piezo1 regulates molecular events, e.g. intracellular calcium (Ca_[i]_) and the activity of FAK. Laser-induced shockwaves have been applied in mechanical destruction of cancer cells; at the same time, it serves as a useful tool to study cancer cell mechanical properties ^16-18^. LIS are produced by the sudden buildup of energy at the focus of a short-pulsed laser beam ^19^. The subsequent microplasma cavitation, followed by microbubble expansion and contraction, and consequently shear stress, can exert a large mechanical force in the surrounding area ^20^. The force is controllable in magnitude by tuning the laser power and can exert instantaneous pressures at the Mega Pascal range to induce intracellular calcium responses ^21-26^. Therefore, laser-induced cavitation and the consequent shear stress can serve as a mechanical stimulator of Piezo1 on single live cells with high precision in space and time^21, 26-28^.

Genetically encoded biosensors based on FRET have enabled the visualization of signaling events in live cells with high spatiotemporal resolution ^29^. They can be applied at the same time to monitor in live cells the Piezo1 activation and its associated Ca_[i]_ dynamics, as well as the FAK activation, a crucial signaling component that regulates cell adhesion and motility ^30^. In this study, by combining mechanical LIS- or chemical ligand-induced Piezo1 activation and live-cell FRET imaging, we revealed a bimodal pattern of Piezo1-induced action of Ca_[i]_ dynamics (transient vs. sustained) and downstream FAK molecular events (activation vs. inhibition), dependent on the degree of Piezo1 activation. These results advance our understanding of Piezo1 activation and its role in perceiving mechanical and chemical cues and regulating intracellular molecular signals.

## 2. Results

To explore whether LIS-induced mechanical stimulation can activate the Piezo1 channel, HEK 293T cells co-transfected with Piezo1-tdTomato and a CFP/YFP-based FRET calcium biosensor ^31^ were examined in a LIS-FRET imaging system ^22^ (Figure 1). In order to create a shockwave sufficient to mechanically stimulate the target cells ^20^ without inducing cell damage, the laser power was precisely controlled in a range between 160 and 180 μW (Figure S1-S2). In a previous study, we showed that cells subjected to LIS treatment with powers at the level up to 342 μW and at a distance of 70 μm remained healthy and did not uptake PI ^22^, a reliable marker for cell necrosis and membrane portion ^25, 26^. Upon one pulse of LIS stimulation of the Piezo1-expressing cells, calcium influx was immediately detected in cells cultured in calcium-containing HBSS media, but not in HBSS without calcium (Figure 2A-B). In control HEK cells transfected only with a calcium biosensor but not Piezo1, there was no significant calcium influx in HBSS media with or without calcium (Figure 2A-B, Figure S3 and Movie S1). In HBSS media without calcium, there was a minor calcium increase in the Piezo1-transfected cells (Figure S3), possibly due to the ER calcium release since Piezo1 can be expressed in ER ^14^. Intermittent shockwave pulses elicited multiple calcium waves (Figure 2C), verifying a healthy response and a lack of shockwave-induced damage on the target cell.

**Figure 1:**
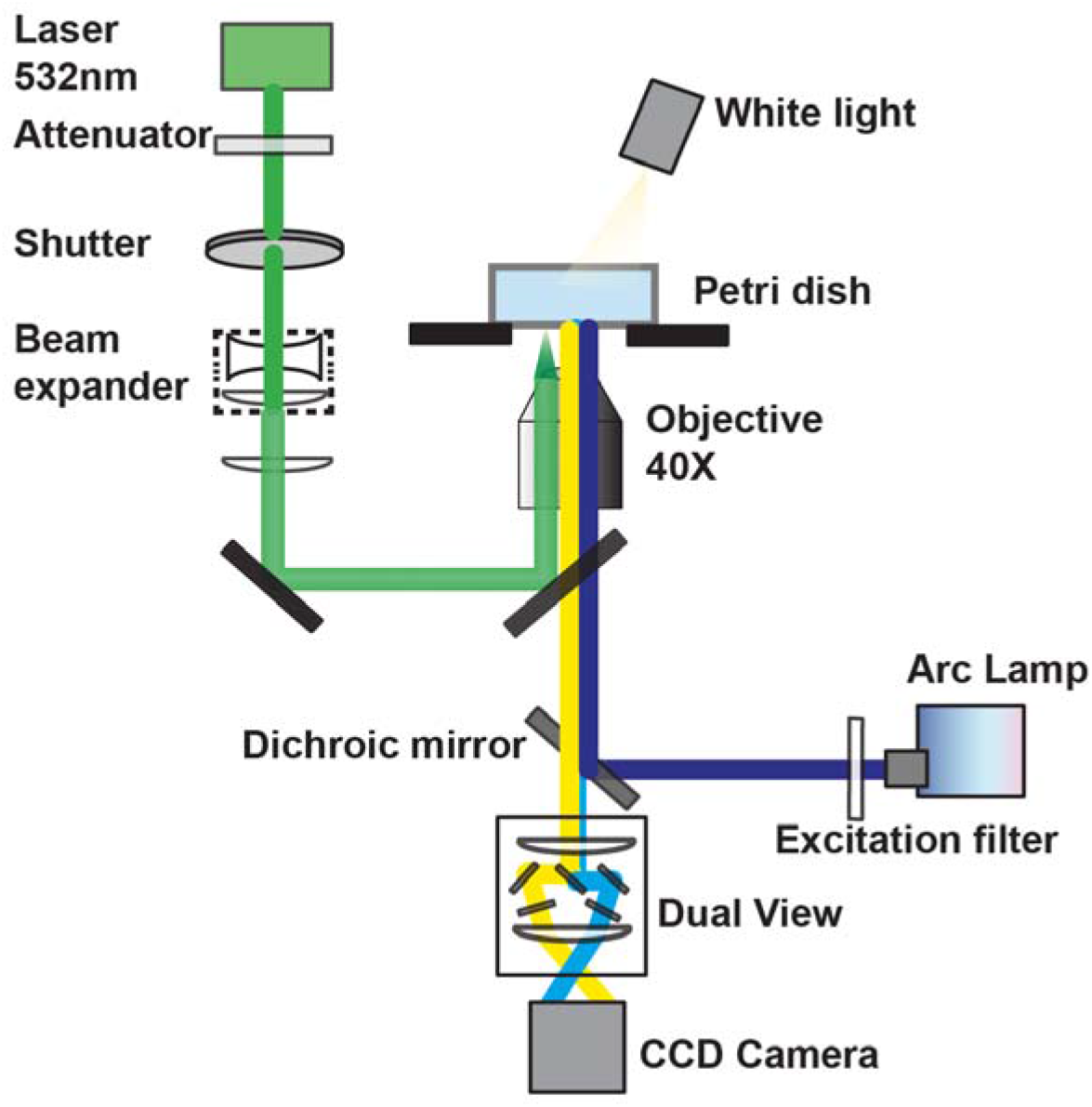
Diagram of an integrated system of Laser-induced Shockwave (LIS) and FRET imaging. The CCD camera captures fluorescent images of FRET biosensors with 480/30 nm and 535/25 nm bandpass filters integrated into the dual view system. The ablation laser is guided through the side port of the microscope. The arc lamp excitation light enters through the backport of the microscope and is filtered with a 440/20 nm bandpass filter.

**Figure 2:**
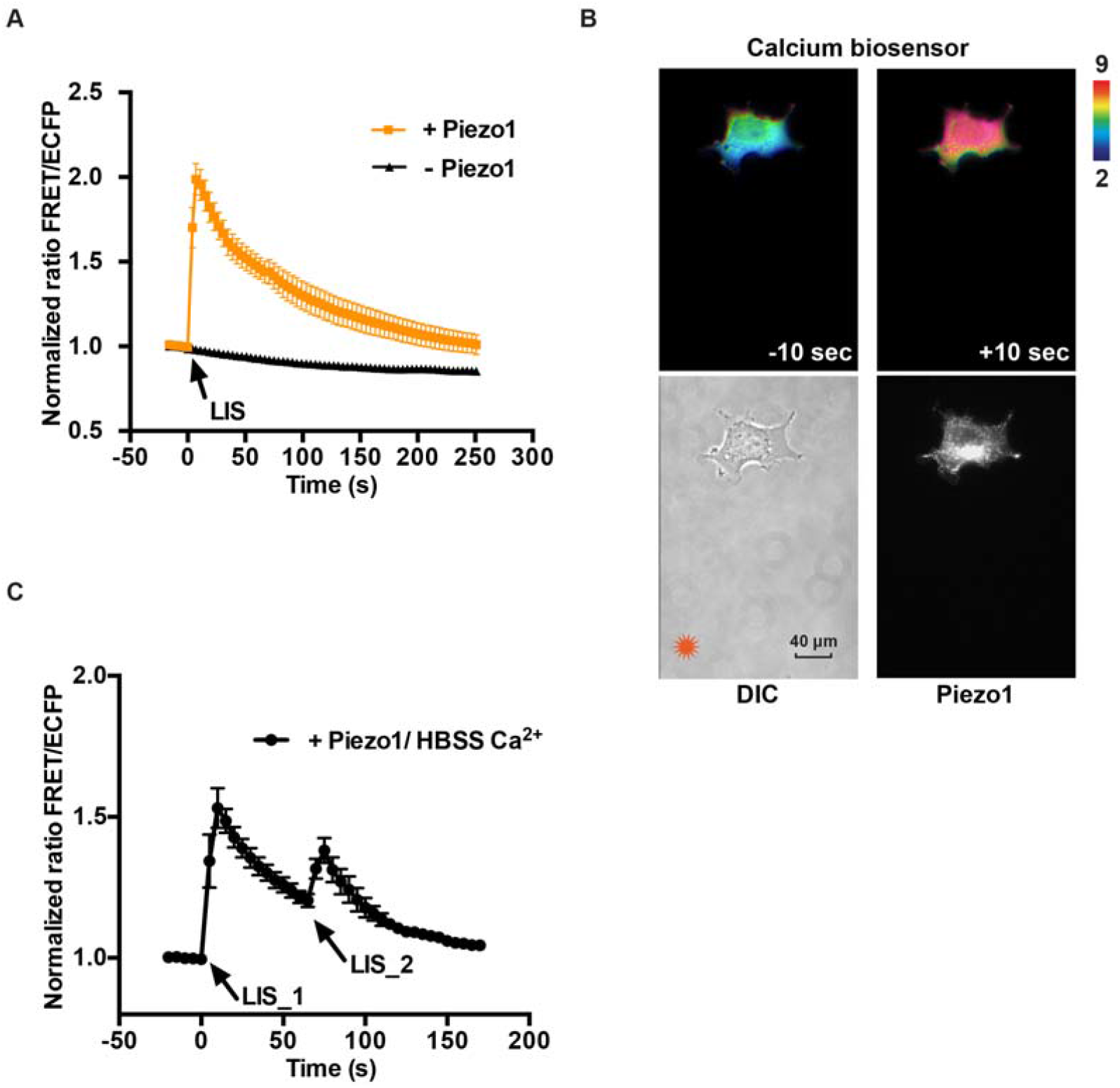
The Piezo1 dependent calcium influx upon LIS stimulation in HEK cells. (**A**) The time courses of normalized FRET ratio (Mean ± SEM) of calcium biosensor before and after shockwave stimulation in HBSS Ca^2+^ medium with (yellow line, n = 6, N = 3) or without the Piezo1 expression (black line, n = 6, N = 3). (**B**) The representative FRET/ECFP ratio images of calcium biosensor in Piezo1-expressed HEK cells before (top left) and after (top right) LIS stimulation. Phase image with laser initiating point (left) and fluorescent image indicating Piezo1 expression (right) were also shown on the bottom panels (Scale bar, 40 μm). Color scale bars in the figures are to show the FRET/ECFP ratios, with cold and hot colors representing low and high ratios, respectively. (**C**) The time course of normalized FRET ratio (Mean ± SEM) of calcium biosensor in the Piezo1-expressing HEK cells before and after multiple shockwave stimulations in HBSS Ca^2+^ medium (n = 6, N = 3). “n” means the total cell number. “N” means the number of individual experiments.

We next examined the effect of this LIS on FAK activity visualized by a FAK FRET biosensor developed earlier by our group ^32^. HEK 293T cells co-transfected with Piezo1-tdTomato and the ECFP/YPet-based FAK FRET biosensor were exposed to LIS in HBSS media containing calcium. One pulse of shockwave stimulation caused an increase of FAK activity in HEK cells transfected with Piezo1, but no significant change in the control HEK cells without Piezo1 expression (Figure 3 and Movie S2). These results indicate that LIS-induced Piezo1 activation could lead to FAK activation, potentially due to the transient calcium influx.

**Figure 3:**
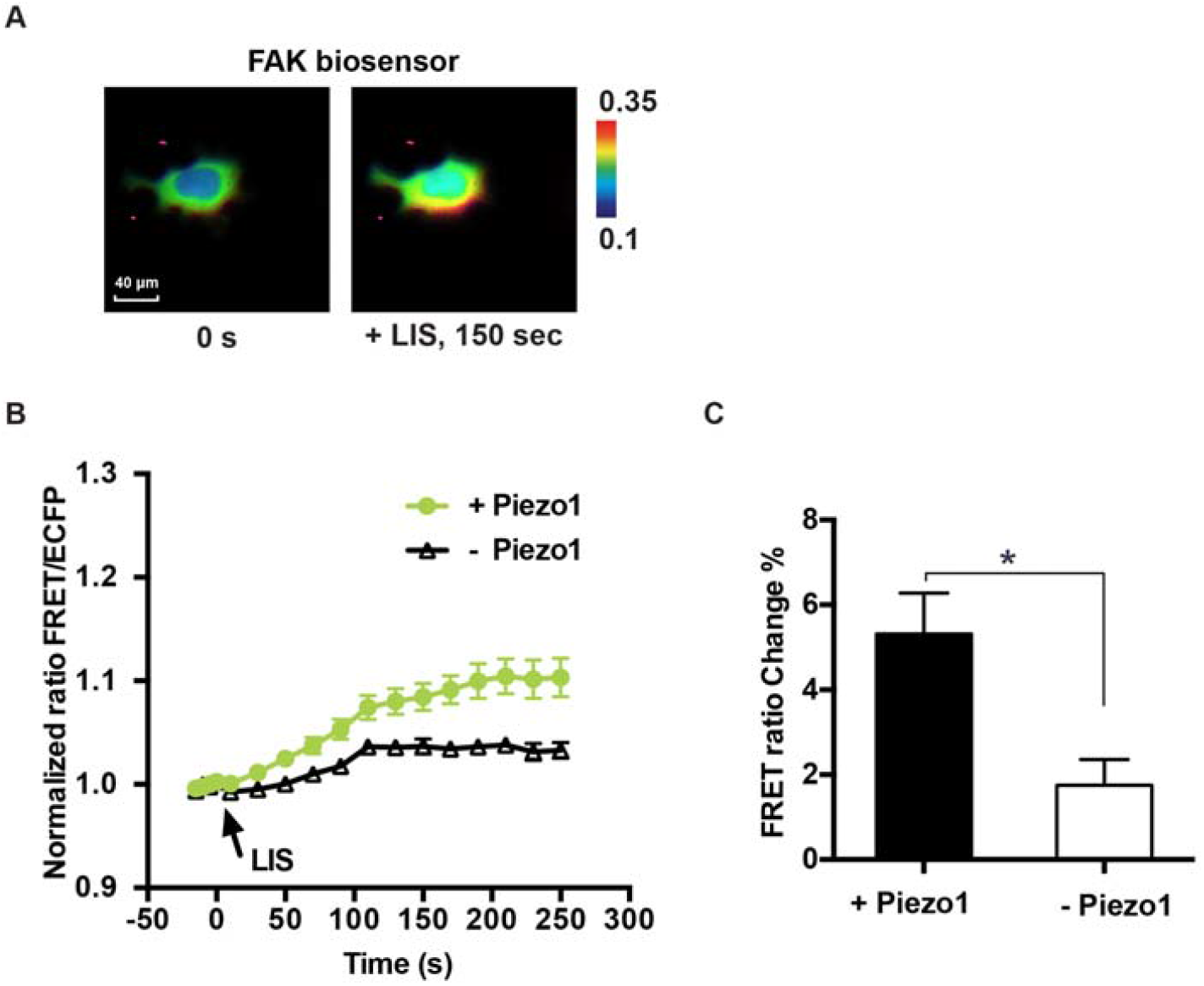
The Piezo1-dependent FAK activation upon LIS stimulation in HEK cells. (**A**) The representative ECFP/FRET ratio images of FAK biosensor in Piezo1 overexpressed HEK cells before (left) and after (right) shockwave stimulation (Scale bar, 40 μm). Color scale bars in the figures are to show the ECFP/FRET ratios, with cold and hot colors representing low and high ratios, respectively. (**B**) The time courses of normalized ECFP/FRET ratio (Mean ± SEM) of FAK FRET biosensor before and after shockwave stimulation in HBSS Ca^2+^ media with (green line, n = 16, N = 3) or without Piezo1 (black line, n = 10, N = 3). (**C**) The percentage change of FRET ratio of the FAK FRET biosensor in HEK cells with or without Piezo1 after shockwave stimulation in HBSS Ca^2+^ medium. *P < 0.05 from student two-tailed t-test. “n” means the total cell number. “N” means the number of individual experiments.

Aside from mechanical stimulation, chemical agonists can also activate the Piezo1 ion channel. To further gain molecular insights on the role of Piezo1 in FAK activation, Yoda1, a synthetic small molecule that has been reported to act as a specific agonist for activating human and mouse Piezo1 ^33^, was applied to stimulate Piezo1. We stimulated HEK cells that have been co-transfected with Piezo1 and the FAK FRET biosensor with 25 μM Yoda1 as a standard dosage ^33, 34^. Surprisingly, Yoda1 stimulation of Piezo1 caused a significant decrease of FAK activity in the HEK cells expressing exogenous Piezo1 (Figure 4A-B); this is opposite to the increase in FAK activity in response to mechanical stimulation of Piezo1 by one pulse of LIS. Hence, it was hypothesized that the LIS-induced mechanical and 25 μM Yoda1-based chemical stimulations result in different modes of the Piezo1 activation. To test this hypothesis, we co-transfected HEK cells with Piezo1 and the calcium FRET biosensor. Upon Yoda1 stimulation at 25 μM, a sustained calcium influx was observed in the cells transfected with Piezo1 (Figure 4C-D), which is significantly different from the transient Ca_[i]_ dynamics observed in the one-pulse LIS stimulated cells (Figure 2A). We also measured the cell viability by calcein-AM dye after 25 μM Yoda1 stimulation, and we didn’t observe cell damage (Figure S4A), suggesting the sustained Ca^2+^ didn’t affect cell health. To further exclude the influence of reporting artifacts of FRET biosensors, we expressed in cells a soluble non-functional FRET biosensor containing ECFP and YPet pair but with its binding domain mutated and tracked its FRET change after stimulated with 25 μM Yoda1. No significant FRET ratio change was observed (Figure S4B). These results suggest that our biosensor-reported biochemical signals induced by Piezo1 activation are not due to the cell morphology changes nor the reporting artifacts of FRET biosensors. The differential FAK responses may hence be attributed to the differences in Ca[i] dynamic patterns upon the mechanical and chemical stimulations.

**Figure 4:**
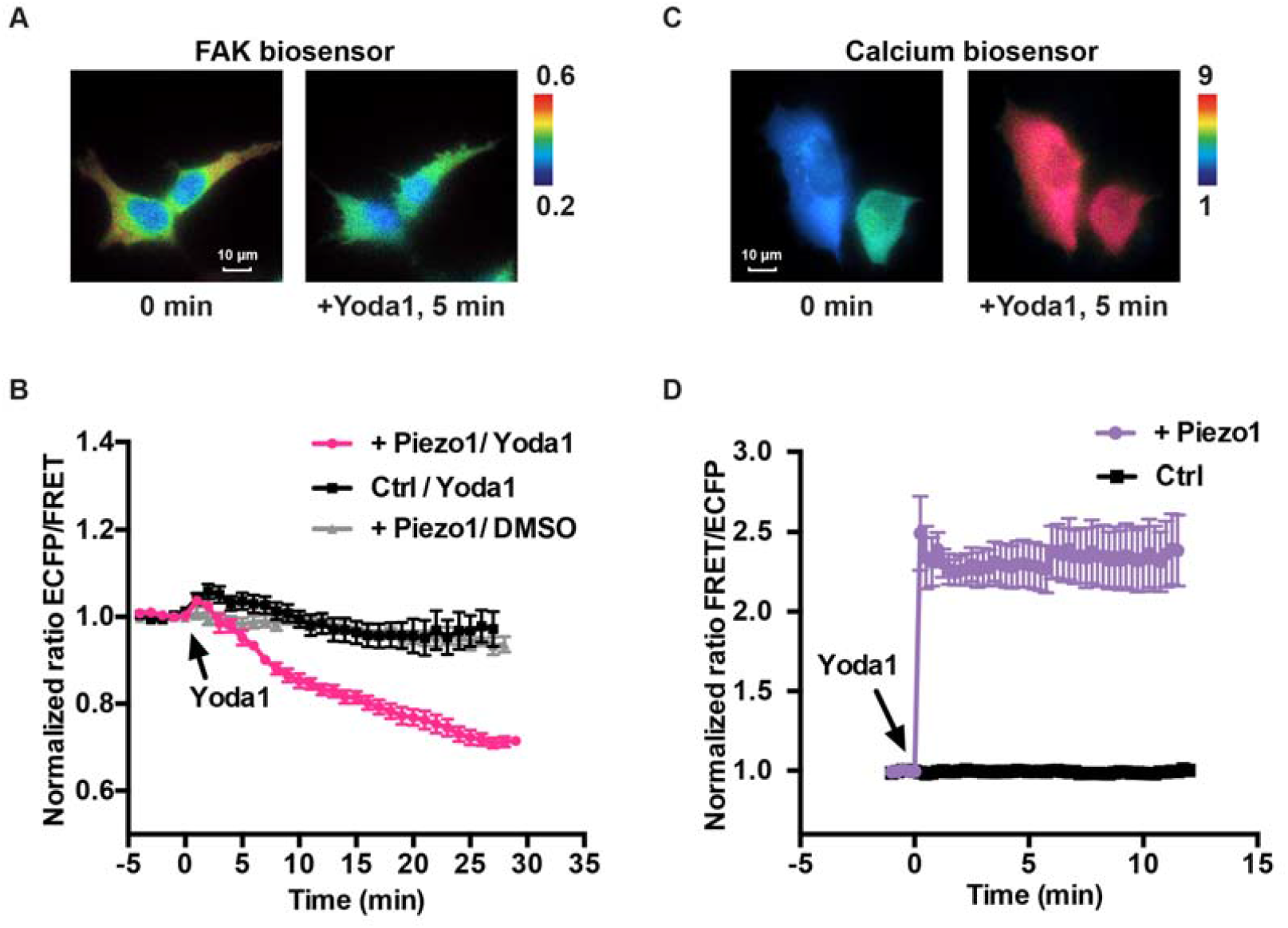
The FAK and calcium activation upon Yoda1 stimulation in the Piezo1-expressing HEK cells. (**A**) The representative ECFP/FRET ratio images of FAK biosensor in Piezo1 expressing HEK cells before and after 25 μM Yoda1 stimulation (Scale bar, 10 μm). Color scale bars in the figures indicate ECFP/FRET ratios, with cold and hot colors representing low and high ratios, respectively. (**B**) The time courses of normalized FRET ratio (Mean ± SEM) of FAK FRET biosensor before and after Yoda1 stimulation with (pink line, n = 10, N = 3) or without Piezo1 (black line, n = 11, N = 3). The grey line represents the DMSO control group (n = 8, N = 3). (**C**) The representative FRET/ECFP ratio images of the calcium biosensor in the Piezo1-expressing HEK cells before and after 25 μM Yoda1 stimulation (Scale bar, 10 μm). Color scale bars in the figures indicate FRET/ECFP ratios, with cold and hot colors representing low and high ratios, respectively. (**D**) The time courses of normalized FRET ratio (Mean ± SEM) of calcium biosensor in the Piezo1-expressing HEK cells (purple line, n = 9, N = 3) before and after 25 μM Yoda1 stimulation in culture medium. “n” means the total cell number. “N” means the number of individual experiments.

To further examine this hypothesis, we titrated the concentration of Yoda1 used to stimulate cells transfected with Piezo1. We observed that a decrease of Yoda1 concentration from 25 to 0.5 μM caused a shift in Ca[i] dynamics pattern from a sustained increase to a transient rise (Figure 5A-B). Correspondently, the FAK response to Yoda1 stimulation changed from a decrease to an increase when the Yoda1 dosage was reduced from 25 to 0.5 μM in the Piezo1-expressing cells (Figure 5C-D). Consistent with these results, increases in the Piezo1 expression level in cells under an intermediate dosage of 2 μM Yoda1 stimulation caused a shift in the dynamic pattern of Ca_[i]_ signaling from transient to sustained (Figure 5E and Figure S5). We also tested additional concentrations of Yoda1 at higher or lower ranges. Indeed, when we increased Yoda1 concentration from 25 μM to 40 μM, a greater reduction in FAK activity was observed. At a lower range, we didn’t observe a significant change in the degree of FAK activity increase, potentially due to less sensitive responses of FAK activities at low levels (Figure S6). These results suggest that the level of Piezo1 activation upon Yoda1 stimulation may have led to differential dynamic patterns of calcium influx to result in FAK activation vs. inhibition.

**Figure 5:**
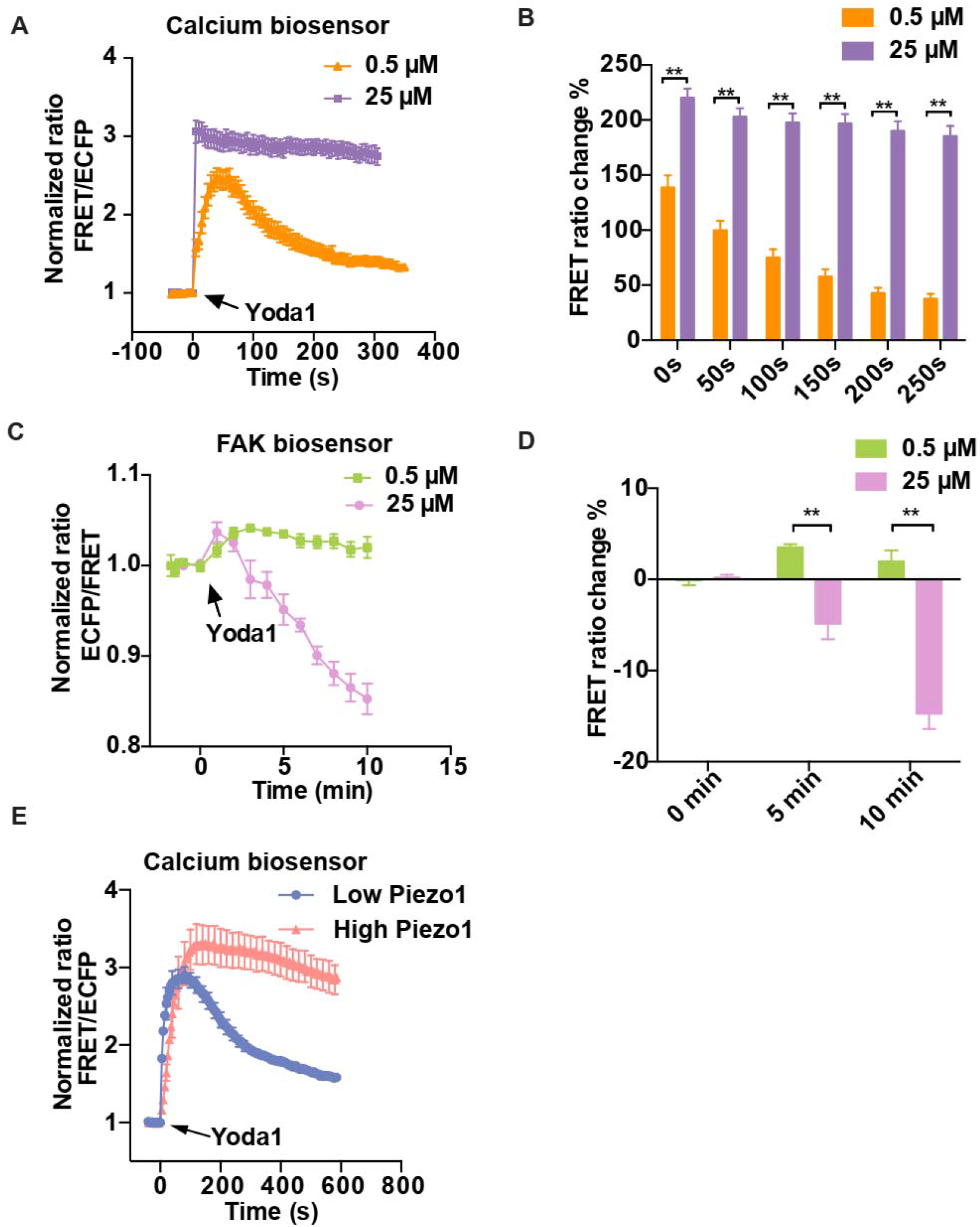
The titration of Yoda1 stimulation in the Piezo1-expressing HEK cells. (**A**) The time courses of normalized FRET ratio of calcium biosensor in the Piezo1-expressing HEK cells upon 0.5 μM (yellow line, n = 10, N = 3) or 25 μM (purple line, n = 8, N = 3) Yoda1 stimulation. (**B**) The percentage changes of FRET ratio decrease of calcium biosensor in Piezo1-expressing HEK cells upon 0.5 μM (yellow bar, n = 8, N = 3) or 25 μM (purple bar, n = 10, N = 3) Yoda1 stimulation at different time points after they reached the peak. \*\**P <* 0.0001 from student two tailed t-test. (**C**) The time courses of normalized FRET ratio (Mean ± SEM) of FAK FRET biosensor before and after 0.5 μM (green line, n = 8, N = 3) or 25 μM (pink line, n = 10, N = 3) Yoda1 stimulation. (**D**) The percentage change of FRET ratio change of FAK biosensor in the Piezo1-expressing HEK cells after 0.5 μM (green bar, n = 8, N = 3) or 25 μM (pink bar, n = 10, N = 3) Yoda1 stimulation at different time points. \*\**P <* 0.0001. (**E**) The time courses of normalized FRET ratio (Mean ± SEM) of calcium biosensor before and after 2 μM Yoda1 stimulation in cells with high (intensity > 4,650 au., orange line, n = 14, N = 3) or low (intensity < 1,000 au., Blue line, n = 5, N = 2) Piezo1 expression.

We thereafter examined whether the bimodal pattern of Peizo1 action observed by chemical stimulation can also be induced by LIS mechanical stimulation with different strength: comparing the effect of multiple vs. single pulse(s) of LIS stimulation. This multiple-pulse LIS stimulation was created by firing laser repeatedly three times within a short time interval of 5 sec (Figure 6A). Multiple LIS stimulation caused a more sustained calcium increase and a decrease of FAK activity in cells transfected with Piezo1, compared to a more transient calcium increase and a slight increase of FAK activity upon single pulsed stimulation (Figure 6B-D). These results are consistent with Piezo1 activation under chemical stimulation, suggesting that different modes of Piezo1 activation can also be triggered by mechanical LIS stimulation. As such, there exists a bimodal pattern of Piezo1 activation-induced Ca_[i]_ dynamics (transient vs. sustained) and downstream FAK (activation vs. inhibition), with the transient mode causing a transient Ca_[i]_ rise and a FAK activation while the sustained mode causing a sustained Ca_[i]_ rise and a FAK suppression (Figure 6E).

**Figure 6.**
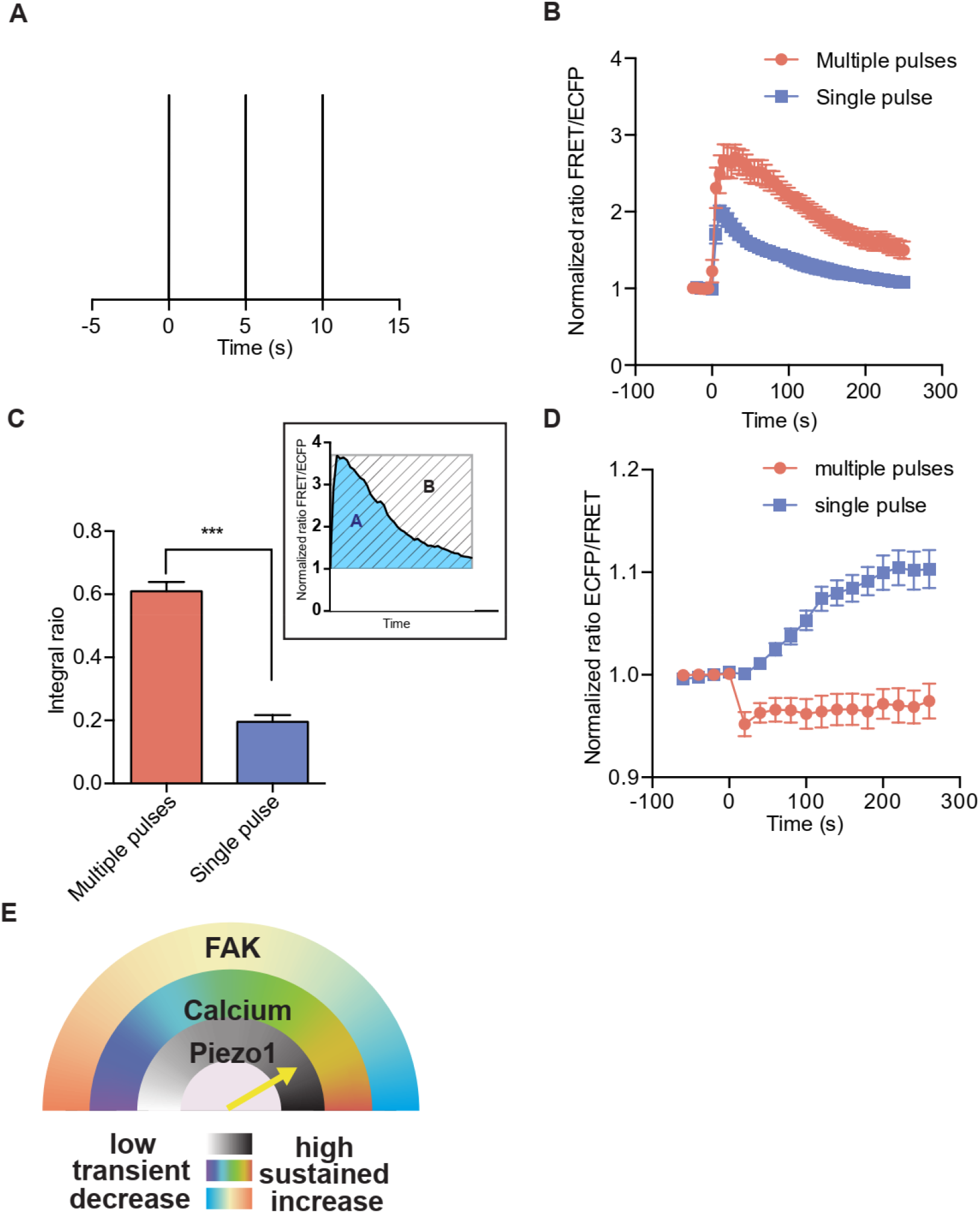
Calcium and FAK activation upon multiple pulsed LIS stimulation in Piezo1 overexpressing HEK cells. (**A**) The stimulation pattern of multiple-pulsed laser stimulation (3 pluses, 5 s interval). (**B**) The time courses of normalized FRET ratio (Mean ± SEM) of d3cpv biosensor before and after three-coupled multiple pulsed or single pulsed LIS stimulation with 5 s interval (blue line, n = 6, N = 3, red line, n = 15, N = 3). **(C**) The percentage change of normalized FRET integral change ratio (See materials and methods, Mean ± SEM) of calcium biosensor in HEK cells with (red bar, n = 15, N = 3) or without (blue bar, n = 12, N = 3) Piezo1 expression before and after multiple-pulsed LIS pulsed stimulations. For each cell, the FRET integral ratio was calculated as the ratio of the area under the time course of the FRET/ECFP ratio (blue area A) to the area under the maximum ratio (patterned area B) over the whole-time course after LIS stimulation. **(D)** The time courses of normalized FRET ratio (Mean ± SEM) of FAK biosensor before and after multiple pulsed LIS stimulation at 5 s interval (blue line, n = 16, red line, n = 36, N = 3). “n” means the total cell number. “N” means the number of individual experiments. (**E**) A cartoon diagram depicting the activation patterns of Piezo1 on downstream calcium (blue or red represents transient or sustained calcium increase pattern, respectively) and FAK signaling (biphasic: blue or red represents a suppression or activation of FAK activity, respectively) upon stimulation. LIS specifically elicits the lower phase (transient mode) of Piezo1 actions (indicated by the pink shade). “n” means the total cell number. “N” means the number of individual experiments.

Next, we investigated the molecular mechanism by which the degree of Piezo1 activation and calcium dynamics lead to the FAK activation or inhibition. The Src-homology-2 (SH2) domain-containing protein tyrosine phosphatase (SHP2), which can be regulated by calcium influx^35^, is a well-known regulator of cell adhesion^36-38^ and FAK activity through its tyrosine phosphatase catalytic domain (PTP domain) directly ^39-41^. SHP2 can also work as an adaptor molecule that binds to phospho-tyrosine containing activators via its two SH2 domains, including insulin receptor substrate 1 (IRS1), GRB2-associated binding protein 1 (GAB1), and non-tyrosine containing substrates such as p53 and fatty acid synthase (FASN)^42-44^. This function of SHP2 is independent of its phosphatase activity ^42-44^. Thus, we hypothesized that SHP2 may work as an intermediate regulator in Piezo1 signaling to regulate FAK differentially in response to different stimulations, through its adaptor or phosphatase function under different conditions. To test this hypothesis and investigate whether and how SHP2 regulates FAK activity downstream of Piezo1 activation and calcium dynamics, we first examined the SHP2 activation pattern following Piezo1 agonist stimulation utilizing a FRET-based SHP2 biosensor^45^. A rapid and sustained SHP2 activation was observed in Piezo1 overexpressing cells after being stimulated with 25 μM Yoda1, while no apparent SHP2 activation in cells stimulated with 0.5 μM Yoda1 (Figure 7A-B). This SHP2 activation is Piezo1 dependent, as no obvious response was observed in Piezo1 negative cells (Figure 7A-B). Such SHP2 activation can lead to the FAK dephosphorylation and suppression (Figure 4B)^39-41^. In contrast, in cells overexpressing a catalytically dead SHP2 (C495S) mutant, FAK was activated when stimulated with 25 μM Yoda1 in cells expressing Piezo1 (Figure 7C), to the same extent as in the control cells stimulated by 0.5 μM Yoda1 stimulation (Figure 7D). Similar results were obtained in cells pretreated with a potent, highly selective SHP2 targeting inhibitor (SHP 099)^46^ that stabilized the auto-inhibited SHP2 conformation and inhibited its catalytic function (Figure 7C, D). These results suggest that SHP2 can lead to FAK activation independent of its catalytic function upon Piezo1 activation. As such, SHP2 may function as a molecular switch to regulate FAK activity in a Piezo1 dependent manner (Figure 7E). SHP2 can be activated under sustained calcium influx to enzymatically dephosphorylate and suppress the FAK kinase; in contrast, under transient calcium, SHP2 serves as an adaptor protein with its enzymatic domain kept in an autoinhibited form to facilitate the activation of FAK.

**Figure 7:**
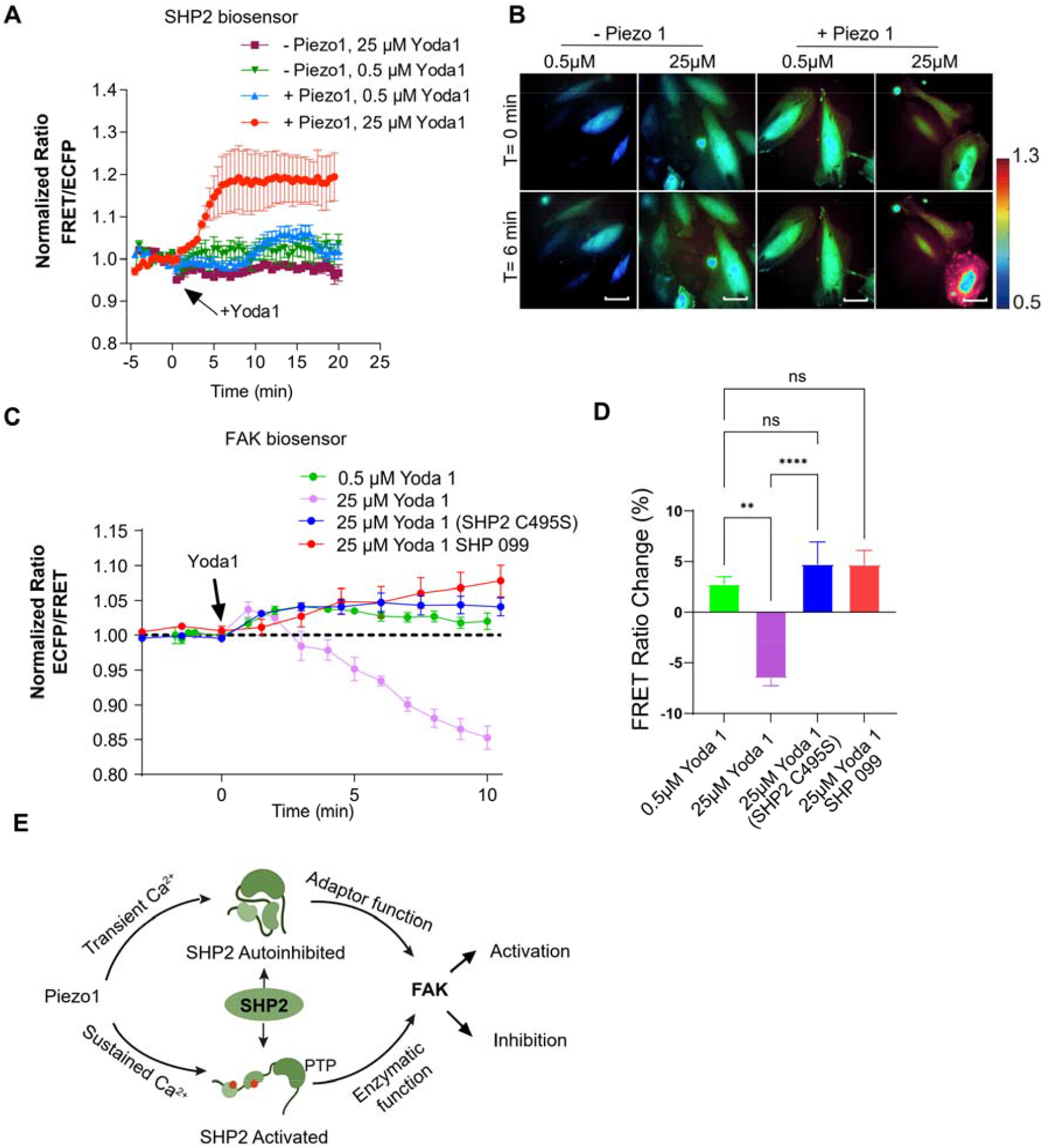
SHP2 works as a molecular switch downstream of Piezo1 to regulate FAK. (A)&(B), The time courses (Mean ± SEM) (A) and representative images of FRET ratio of SHP2 biosensor (Y542F) before and after 25 μM or 0.5 μM Yoda1 stimulation in cells with or without Piezo1 overexpression (red bar, n = 5, N=3; blue bar, n = 5, N=3; Purple bar, n = 11, N=3; green bar, n = 5, N=3). “n” means the total cell number. “N” means the number of individual experiments. (C). The time courses of normalized FRET ratio (Mean ± SEM) of FAK FRET biosensor under different treatments. (From Top to the bottom, n=8, 10, 14 and 14 respectively; SHP 099, the cells were pretreated with 10 μM SHP099; SHP2 C495S, the cells overexpressing SHP2 protein with a non-functional catalytic domain). (D). The percentage change of FRET ratio changes of FAK biosensor in the Piezo1-expressing cells in C at 5 minutes. (From left to right, n=8, 10, 14 and 14 respectively; one-way ANOVA, **P=0.006, ****P<0.0001, NS=not significant.) (E). Proposed working model of FAK regulation by Piezo1 in which SHP2 works as a molecular switch.

## 3 Discussion

In this study, we revealed a bimodal pattern of Piezo1 activation-induced Ca_[i]_ dynamics (transient vs. sustained) and downstream FAK (activation vs. inhibition), with the transient mode causing a transient Ca_[i]_ rise and a FAK activation while the sustained mode causing a sustained Ca_[i]_ rise and a FAK suppression. By combining FRET imaging with laser-based photonics, we have successfully monitored the distinct molecular signaling events with high spatiotemporal resolutions in live cells following the mechanosensor Piezo1 activation by LIS mechanical stimulation and the ligand chemical stimulation.

Interestingly, high levels of Piezo1 activation, either attributed to high dosages of Yoda1 application or to high expressions of Piezo1, caused sustained calcium signaling and FAK suppression. Tuning down the Piezo1 activation level can lead to a switch of calcium and FAK responses, with the dynamic Ca_[i]_ signaling becoming more transient, accompanied by an activation of FAK. This is consistent with previous reports that transient calcium influx could lead to increased FAK phosphorylation ^47, 48^. These results indicate that the strength of Piezo1 activation can lead to two distinctive activation patterns (bimodal) of downstream signaling, with low levels of Piezo1 activation triggering transient calcium signaling and FAK activation (transient mode) while high levels of Piezo1 activation led to sustained calcium signaling and FAK suppression (sustained mode) (Figure 6E). We believe that calcium plays a central role in the bimodal actions of FAK. It is well established that the kinetics and amplitude of Ca^2+^ signals allow the activation of specific Ca^2+^ targets and potentially the differential focal adhesion (FA) characteristics^49^. Specifically, the process of cellular adhesion, involves the dynamic regulation of FAs complexes ^50, 51^. FAK can phosphorylate local substrate molecules and create docking sites for the molecular recruitment and further assembly of the FA complex ^52, 53^. We indeed examined FAK dynamics by tracking a focal adhesion-associated adaptor protein paxillin^54^. We stimulated Piezo1-expressing HEK cells with 25 μM or 0.5 μM Yoda1, and observed a decrease of paxillin intensity in 25 μM Yoda1 treated cells, but a slight increase of paxillin intensity with 0.5 μM Yoda1 treatment (Figure S6A-D). These observations are also consistent with the FAK suppression and activation caused by 25 μM and 0.5 μM Yoda1, respectively.

Further investigations suggest that SHP2 may serve as a molecular switch downstream of Piezo1 to regulate FAK activity, which leads to different FAK responses upon the different levels of Piezo1 activation. SHP2 can be activated under sustained calcium influx to enzymatically dephosphorylate and suppress the FAK kinase, while serving as an adaptor protein independent of its enzymatic function to facilitate the activation of FAK under transient calcium. While further investigations will be warranted to reveal more molecular insights, this study should shed new light on how the mechanosensitive Piezo1 regulates cell adhesion and paves the way to a better understanding of Piezo1 related mechanotransduction processes.

Our findings of the bimodality of Piezo1-induced signaling should also help to advance our understanding of the opposite effects of Piezo1 knockdown on the migration of two cell lines: a decrease for gastric tumor cell ^55^, but an increase for lung epithelial cells ^14^. Since stomach and lung cells have a low and high level of Piezo1 expression, respectively^10^, the knockdown of Piezo1 on these cells may have tuned their Piezo1 levels and hence sensing strengths differentially in response to environmental cues. This may ultimately lead to the different outcomes in perturbing cell migration, in which FAK plays a crucial role. As such, our results provide some insights into the paradox that the same manipulation of Piezo1 under different cell types can lead to differential outcomes in cell migration ^13, 56^.

### Experimental Section

#### DNA plasmid

The construct of the mPiezo1-tdTomato was obtained from Dr. Patapoutian’s lab ^12^. The D3cpv calcium FRET biosensor was previously described ^31^, the cytosolic FAK FRET biosensor was previously developed and described ^32^.

#### Cell culture and transfection

HEK 293T cell line was a kind gift from Dr. Patapoutian’s lab ^7^. Cell culture reagents were obtained from Invitrogen. Cells were maintained in Dulbecco’s modified Eagle medium (DMEM) supplemented with 10% fetal bovine serum (FBS), 2 mM L-glutamine, 1 unit/ml penicillin, 100 μg/ml streptomycin, and 1 mM sodium pyruvate. Cells were cultured in a humidified 95% air, 5% CO_2_ incubator at 37 °C. FUGENE 6 (Invitrogen) was used for transfection of DNA plasmids. Cells were tested 36-48 hr post-transfection.

During imaging, cells were cultured in fibronectin-coated cover-glass-bottom dishes and maintained in Hanks Buffered Saline solution (HBSS) with or without Calcium and Magnesium (Invitrogen, Carlsbad, CA). Chemical stimulant, Yoda1, was purchased from Thermo fisher scientific. Calcein AM was purchased from Thermo fisher scientific. Cells were stained with Calcein AM according to manufacturer’s manual.

#### Laser-Shockwave System

The laser used for the shockwave stimulation is a 532 nm wavelength, 100 Hz repetition rate, 2 ns pulse width Coherent Flare system (Spectra-Physics). Laser power is attenuated by rotating an optical polarizer mounted in a stepper-motor-controlled rotating mount (New Port). A mechanical shutter (Vincent Associates) with a 10-15 ms duty cycle is gated to allow 1-2 laser pulses to pass through when firing. The beam expanders are used to adjust the beam diameter of the lasers to fill the back aperture of the objective. The custom laser entry port (CLEP) is placed underneath the objective (phase III, 40x, NA 1.3, oil immersion) of a Zeiss Axiovert 200M microscope. The filter on the CLEP was chosen to direct the laser lights onto the lower right corner of the image screen.

#### Laser-induced shockwave (LIS)

In order to understand precisely the forces presented in the shockwave system, we followed a similar method in a previous publication to measure and calculate forces ^20^. The time resolved shockwave signal was determined by monitoring the size of each bubble created by the laser focus plasma. In order to achieve this, a low power measurement laser at 633 nm was arranged to be coaxial with the shockwave beam. The transmitted power of the beam was monitored using a fast photodiode. Laser focus was at 10 μm above the substrate. Upon the creation of a bubble during the measurement, laser was deflected and the signal was captured.

A series of measurements at different powers between time-averaged 160 μW and 293 μW measured at the back aperture were conducted. The peak beam deflection and the time of bubble rise and collapse were recorded for calculation.

By utilizing the Rayleigh Equation for bubble growth:

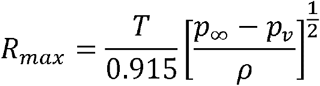

Where *R*_max_ is the maximum bubble radius, T is the time for bubble collapse.

*P*_∞_is the atmospheric pressure, *P*_*v*_ is the vapor pressure within the bubble and *P* is the density of the fluid. This information can then be fit into the Gilmore Equation and applied to a Logarithmic fit function to calculate the time dependent bubble expansions.

We calculated the radial shear forces in our system by applying the methods shown previously ^19, 22, 57^

The radial distribution of shear forces was shown below.

| *Time averaged power $\mu\text{W}$* | *Optical intensity $(\text{GW}/\text{m}^2)$* | *Pulse Energy $(\mu\text{J})$* | *Bubble Energy $(\mu\text{J})$* | *Peak Shear Stress $(\text{MPa})$* |
| --- | --- | --- | --- | --- |
| $164 \pm 0.1$ | $1.24\text{e}+4 \pm 1\text{e}+3$ | $1.6 \pm 0.04$ | $0.21 \pm 0.02$ | $21.10 \pm 4$ |
| $170 \pm 0.1$ | $1.28\text{e}+4 \pm 1\text{e}+3$ | $1.7 \pm 0.04$ | $0.22 \pm 0.02$ | $21.38 \pm 4$ |
| $180 \pm 0.1$ | $1.36\text{e}+4 \pm 1\text{e}+3$ | $1.8 \pm 0.04$ | $0.23 \pm 0.02$ | $21.84 \pm 4$ |

We noted that cells have been shown to survive through pressures up to 120 MPa ^58^, as cell lysis is not only dependent on the peak shear but also on the duration of the applied shear stress, as well as cell types and bubble expansion radius^28^. The cell lysis was observed to occur at distances below 120 μm away from the epicenter of the cavitation with our current experimental configuration. In our mechano-stimulation experiments, cells were positioned more than 190 μm away from the cavitation epicenter to avoid cell damages.

#### Image acquisition

Images of shockwave stimulation were collected by ORCA-Flash4.0 V2 Digital CMOS camera (Hamamatsu) passing a 440DF20 excitation filter, a 455DRLP dichroic mirror, and two emission filters placed in a dual view adaptor (480DF30 for ECFP and 535DF25 for EYFP). Images of agonist stimulation were collected from a Nikon microscope with a charge-coupled device (CCD) camera with the same filter setting. For FAK biosensor, the pixel-by-pixel ratio images of ECFP/FRET were calculated based on the background-subtracted fluorescence intensity images of CFP and YFP by the Metafluor program to allow quantification and statistical analysis of FRET responses. For calcium biosensor, the emission ratio of FRET/ECFP was used to represent calcium activity. For each cell, the FRET integral ratio was calculated as the ratio of the area under the time course of the FRET/ECFP ratio (blue area A) to the area under the maximum ratio (patterned area B) over the whole time course after LIS stimulation. The emission ratio images were shown in the intensity modified display (IMD) mode. In terms of emission ratios quantifying the biosensor activations, we used ECFP/FRET for FAK biosensor, and FRET/ECFP for calcium biosensor. The reason is that the two biosensors are designed with different FRET activation mechanisms. For FAK biosensor, FAK activation causes a decrease in FRET efficiency, while for calcium biosensor, calcium activation causes an increase in FRET efficiency. Therefore, we used the indicated emission ratios described above to represent biosensor activations, so that an increase in the emission ratio indicates an activity increase in the target molecule.

#### Focal adhesion analysis

The detection method of FA locations based on the GFP fluorescence intensity (FI) images was as previously described ^59^. In brief, the image analysis for focal adhesion was conducted using our customized software fluocell developed in MATLAB. The source code of *fluocell* is published at Github (https://github.com/lu6007/fluocell). All the images were background-subtracted and smoothed using a median-filter with a window size of 3 × 3 pixels. The GFP-paxillin images were used to compute the masks of the cells. The cell masks were then divided into four layers with the outer layer representing the lamellipodium region. After that, a fan-shaped region was selected in the first image of the video sequence to identify the location of active lamellipodia (Figure S6A and S6B). The interception of this fan region and the outer layer of the cell mask were utilized for all the images in the sequence to represent the active lamellipodium region of interest (ROI). As such, the paxillin-GFP images were processed with high-pass filter which removed diffusive cytosolic background (Fig. S6B) and a threshold value was used to detected paxillin-labelled focal adhesions in the filtered image. Total FI of the detected paxillin focal adhesions were quantified in the ROI over time and normalized such that the average value before treatment was 1. The normalized time courses of total FI were then collected for further analysis and statistical comparison.

## Acknowledgements

The authors thank Dr. Ardem Patapoutian^6,7^ for his generosity in providing materials and advice. This work is supported by grants from NIH GM125379, GM126016, CA204704, CA209629, and HL121365 (Y. Wang), NSF CBET1360341, DMS1361421 (Y. Wang and S.L.), the Beckman Laser Institute Foundation, AFOSR FA9550-08-1-0284, Air Force Office of Scientific Research under award number FA9550-17-1-0193, Beckman Laser Institute Inc. Foundation (M.W.B.). The funding agencies had no role in study design, data collection and analysis, decision to publish, or preparation of the manuscript.

## Author Contributions

Y.P., S.C., M.W.B., L.L., and Y. W. designed research; Y.P., L.L., L.Z.S., D.P., C.Y., C.C., and V.G. performed research; S.W. contributed new reagents; Y.P. and S.L. analyzed data; Y.P., D.P., S.C., M.W.B., L.L., and Y.W. wrote the paper.

