## Supplementary figures for "Mechanosensor Piezo1 Mediates Bimodal Patterns of Intracellular Signaling"

[1]Department of Bioengineering, [2]Institute of Engineering in Medicine, [3]Department of NanoEngineering, [4]Department of NanoEngineering, University of California, San Diego, La Jolla, CA 92093, USA, [5]Beckman Laser Institute and Medical Clinic, University of California, Irvine, Irvine, CA 92612, Department of Cell Biology, [6]Dorris Neuroscience Center, The Scripps Research Institute, La Jolla, CA 92037, USA, [7]Genomic Institute of the Novartis Research Foundation, San Diego, CA 92121, USA

[*] To whom correspondence should be addressed:

Yingxiao Wang, Ph. D.

Shu Chien, Ph.D., M.D.

Michael Berns, Ph.D.

Longwei Liu, Ph. D.

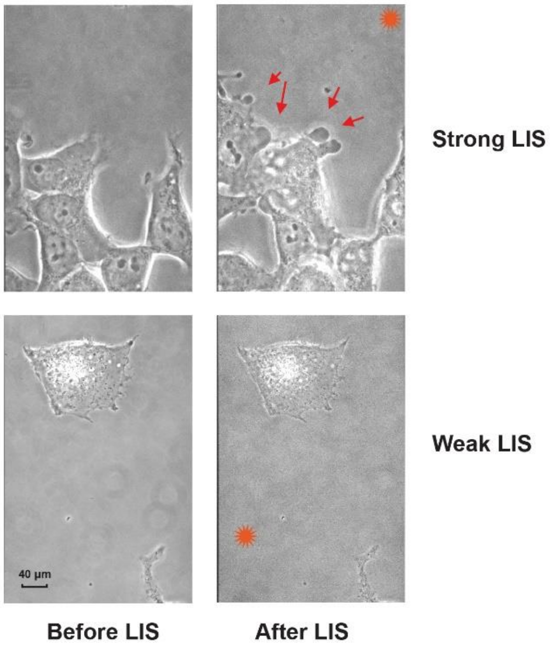


**Figure S1**

Shockwave produced by strong laser power can cause cell damage. The representative phase image of HEK cells before (top left) and after (top right) strong shockwave (200 μW) stimulation. Cells were damaged upon stimulation, as the bubbling indicated by red arrows. The representative DIC images of HEK cells before (bottom left) and after (bottom right) weak shockwave (180 μW) stimulation were shown to indicate that cells were alive and not damaged upon stimulation.

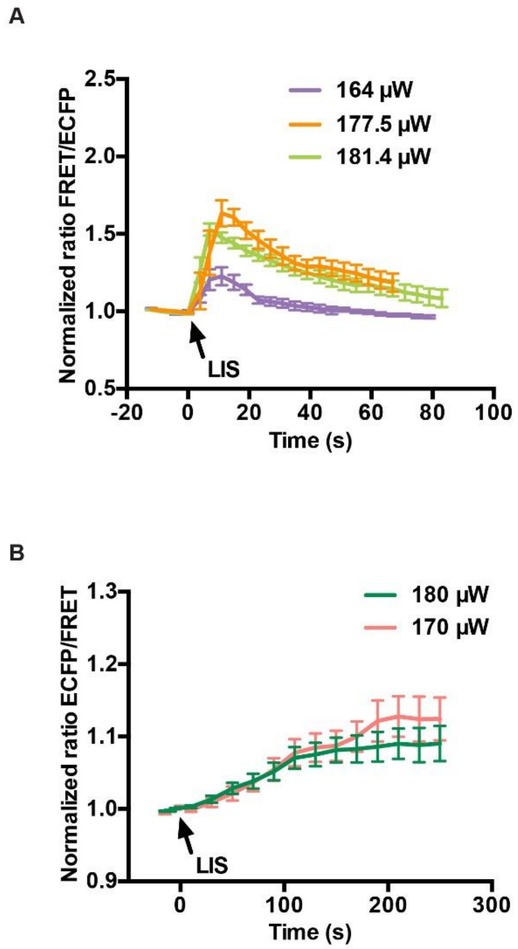


**Figure S2**

Calcium and FAK activation upon LIS stimulation in Piezo1 overexpressing HEK cells with different shockwave strengths. (**A**) The time course of normalized FRET ratio (Mean ± SEM) of d3cpv biosensor before and after LIS stimulation at different laser power strengths (purple line, n = 4, yellow line, n = 7, green line, n = 5, N = 3). (**B**) The time course of normalized FRET ratio (Mean ± SEM) of FAK biosensor before and after LIS stimulation at different laser power strengths (pink line, n = 8, green line, n = 8, N = 3). “n” means the total cell number. “N” means the number of individual experiments.

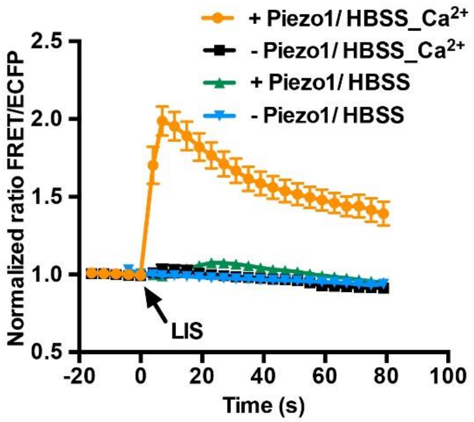


**Figure S3**

The time courses of normalized FRET/CFP ratio (Mean ± SEM) of d3cpv calcium FRET biosensor before and after shockwave stimulation in HBSS $Ca^{2+}$ media with (orange line, n = 6, N = 3) or without Piezo1 (black line, n = 6, N = 3), or in HBSS only media with (green line, n = 18, N = 3) or without Piezo1(blue line, n = 14, N = 3). “n” means the total cell number. “N” means the number of individual experiments.

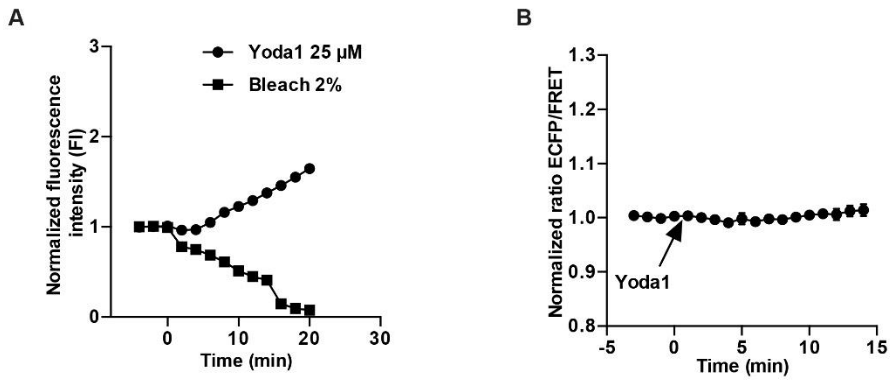

**Figure S4**

Cell viability and response upon Yoda1 stimulation in Piezo1 overexpressing HEK cells. (**A**) The time courses of normalized fluorescence intensity (FI) of calcein-AM (Mean ± SEM) before and after 25 μM Yoda1 (circle, n = 56, N = 3) or 2% bleach (square, n = 39, N = 3) stimulation. The quantified time course of total fluorescent intensity (FI, EX470 nm/ EM525 nm) was normalized such that the average value before treatment was 1. (**B**) The time courses of normalized FRET ratio (Mean ± SEM) of a soluble FRET biosensor (Mean ± SEM) before and after 25 μM Yoda1 stimulation (n = 19, N = 3). “n” means the total cell number. “N” means the number of individual experiments.

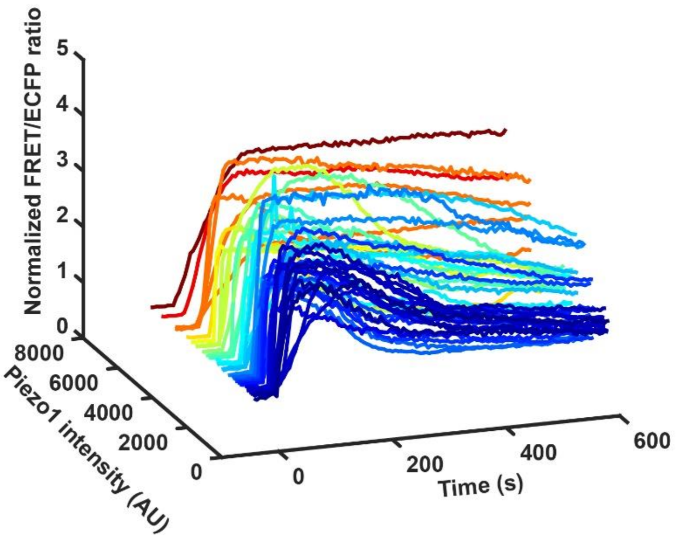


**Figure S5**

Piezo1 level dependent calcium influx upon Yoda1 stimulation in HEK cells. The time courses of normalized FRET ratio (Mean ± SEM) of d3cpv biosensor before and after

2 μM Yoda1 stimulation in HEK cells expressing different levels of Piezo1 (n = 44, N = 3). "n" means the total cell number. "N" means the number of individual experiments.

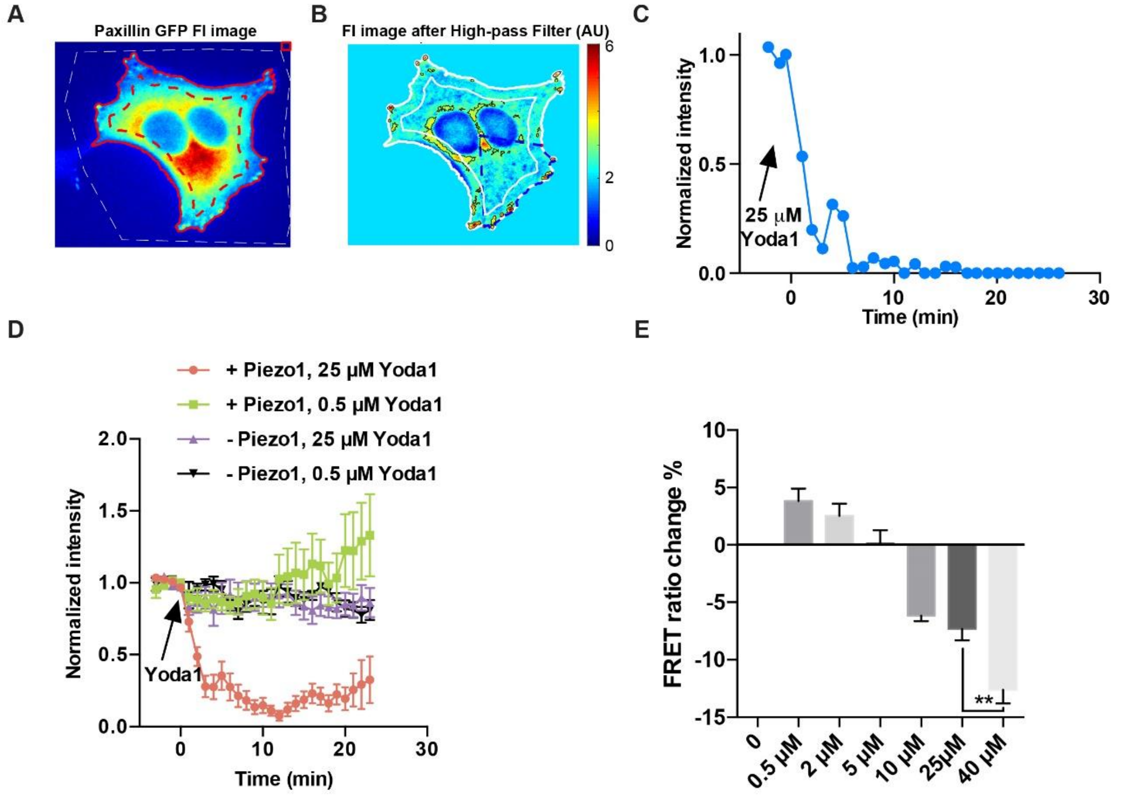


**Figure S6**

The Focal adhesion and FAK responses upon Yoda1 stimulation in the Piezo1-expressing HEK cells. (**A**) shows the GFP FI image of a cell co-expressing GFP-paxillin and Piezo1 in pseudo color. The detected cell body is outlined in red (solid line), the outer layer corresponding to lamellipodia lies between the solid red and dashed red lines. A mask outlined in dashed white was used to exclude unspecific objects. The red rectangle at the top right corner indicates the background region used for preprocessing. (**B**) shows the GFP FI image after the high-pass filter was applied to remove diffusive cytosolic background. The lamellipodia region is outlined between two solid white lines. A fanshaped region (dashed blue line) was selected to identify polarized lamellipodia. The detected focal adhesions with high FI were outlined in black. The total intensity of paxillin-GFP in the interception of detected FA, selected fan, and lamellipodia was quantified over time. (**C**) The quantified time course of total GFP fluorescent intensity (FI) was normalized such that the average value before treatment was 1. The normalized time course for the cell shown in (A) is plotted. (**D**) The time courses of normalized GFP fluorescent intensity (FI) before and after 25 μM or 0.5 μM Yoda1 stimulation in cells with or without Piezo1 overexpression (red bar, n = 8, N=3; green bar, n = 13, N=3; purple bar, n = 7, N=3; black bar, n = 5, N=3). (**E**) The percentage change (at 5 min) of FRET ratio of the FAK FRET biosensor in the Piezo1-expressing HEK cells after different concentrations of Yoda1 stimulation (0.5 μM Yoda1, n = 31, 2 μM Yoda1, n = 29, 5 μM Yoda1, n = 51, 10 μM Yoda1, n = 42, 25 μM Yoda1, n = 21, 40 μM Yoda1, n = 21, N = 3). **P < 0.005 from student two-tailed t-test. "n" means the total cell number. "N" means the number of individual experiments.

**Supplementary Movies legends:**

**Movie S1** The representative FRET/ECFP ratio movie of d3cpv biosensor in Piezo1 overexpressed HEK cells before and after shockwave stimulation. Laser initiating point was at the frame where "laser on" was labeled.

**Movie S2** The representative ECFP/FRET ratio movie of FAK biosensor in Piezo1 overexpressed HEK cells before and after shockwave stimulation. Laser initiating point was at the frame where "laser on" was labeled.
